# A Multicenter Preclinical MRI Study: Definition of Rat Brain Relaxometry Reference Maps

**DOI:** 10.1101/2020.02.06.928549

**Authors:** Tristan Deruelle, Frank Kober, Adriana Perles-Barbacaru, Thierry Delzescaux, Vincent Noblet, Emmanuel L. Barbier, Michel Dojat

**Author notes:** Correspondence should be addressed to: Michel Dojat, Grenoble Institut des Neurosciences - Inserm U1216, Site Santé, Chemin F. Ferrini, 38700 La Tronche, France.

## Abstract

Similarly to human population imaging, there are several well-founded motivations for animal population imaging, the most notable being the improvement of the validity of statistical results by pooling a sufficient number of animal data provided by different imaging centers. In this paper, we demonstrate the feasibility of such a multicenter animal study, sharing raw data from forty rats and processing pipelines between four imaging centers. As specific use case, we considered the estimation of T1 and T2 maps for the healthy rat brain at 7T. We quantitatively report about the variability observed across two data provider centers and evaluate the influence of image processing steps on the final maps, by using three fitting algorithms from three centers. Finally, to derive relaxation time values per brain area, two multi-atlas segmentation pipelines from different centers were executed on two different platforms. In this study, the impact of the acquisition was 2.21% (not significant) and 9.52% on T1 and T2 estimates while the impact of the data processing pipeline was not significant (1.04% and 3.33%, respectively). In addition, the computed normality values can serve as relaxometry reference maps to explore differences to animal models of pathologies.

## INTRODUCTION

In the clinical domain, multicenter studies are common. Their main objective is to gather sufficient patient data in a reasonable period of time, and to improve the statistical power and consequently the robustness of the reported results. Multicenter studies also set the basis for developing and validating quantitative and reproducible imaging biomarkers. Although the animal/preclinical imaging community faces the same challenges regarding the need for combining larger sets of data, to our knowledge, no population study across multiple centers has been reported so far. Consequently, only few tools are available to facilitate preclinical data pooling. Moreover, there is a clear lack of large actions for standardization of image acquisition conditions and post-processing techniques. Finally, there are no reliable commonly adopted preclinical imaging biomarkers for differentiating normal vs pathological conditions.

The aim of the present work was to assess the feasibility of multicenter preclinical studies. We thus considered a specific use case: quantitative mapping of T1 and T2 in the healthy rat brain at 7T. To facilitate pooling of the preclinical MR datasets, we used the SHAring NeurOImaging Resources (Shanoir) environment, which was initially developed for the web-oriented management of collaborative neuroimaging projects in Humans (Barillot et al., 2016), but recently extended for preclinical studies (Small Animal Shanoir, SAS). We quantitatively report about the variability observed across data provider centers and evaluate the impact of different image processing pipelines. We finally discuss the feasibility of small animal population studies. To promote data sharing in the preclinical domain, both raw and processed datasets as well as the employed processing pipelines have been made available (see Discussion Section for details).

## MATERIALS AND METHODS

### Distribution of tasks between centers

Two centers, GIN (C1) and CRMBM (C2) hosted the animals and performed MRI brain acquisitions. Three centers (C2), MIRCen (C3) and ICube (C4) provided the processing pipelines.

### Animals

Twenty Sprague-Dawley rats (male, Janvier Labs, Paris France, mean weight 279±40 g ([min: 249.5g, max: 314g], details in SM Table 1) were scanned in two imaging centers (C1 and C2) at the age of 7 weeks (except for C1 where half of the rats were scanned at an age of 10 weeks). Animals were anaesthetized with isoflurane (2% in air at C2 and 2% in a mixture of Air and O_2_ (7:3) at C1) that was delivered via a nose cone during the experiment. Animals were positioned in prone position on an animal bed (Bruker Biospin, Ettlingen, Germany). Breath rate was monitored using a pneumatic pillow sensor placed under the abdomen. Body temperature was measured with a rectal probe and maintained in the normal range at 36.2±1.4 °C using a heated blanket. To evaluate acquisition reproducibility, three rats were scanned twice, one rat (Subject 32) at C2 at a 2-day interval and two rats (Subject 21 and Subject 22) at C1 at a 3-day interval. All experiments were approved by the local ethics committee of each center and were in full compliance with the guidelines of the European Union (EUVD 86/609/ EEC) for the care and use of the laboratory animals. Experiments were performed under permits from the French Ministry of Agriculture (nº 380945 and A3851610008 for experimental and animal care facilities for C1 and G130555 for C2).

**Table 1:** Equipment characteristics, Fitting and Segmentation methods. Where A, B, T1 and T2 are the parameters to be estimated. See text for details.

| **Acquisition** |  | **GIN (aC1)** | **CRMBM (aC2)** |
| --- | --- | --- | --- |
| Scanner Model |  | Bruker BioSpin MRI GmbH –<br>Biospec 70/20 7.0T | Bruker BioSpin MRI GmbH –<br>Pharmascan B-C 70/16US 7.0T |
| Transmitting coil |  | Linear Volumetric Coil 72 mm | Quadrature Resonator 72 mm |
| Receiving coil |  | Rat Head Coil (Surface) 1 channel | GIN coil |
| Gradient system |  | BGA12S2 (110mm, max 660 mT/m,<br>slew rate 4570 T/m/s) | BGA 9SHP (90mm, max 750 mT/m,<br>slew rate 6840 T/m/s) |

| **Fitting** |  | **GIN (fC1)** | **CRMBM (fC2)** | **MIRCen (fC3)** |
| --- | --- | --- | --- | --- |
| Software |  | MATLAB R2015a | ImageJ v1.51 | BrainVISA/Python |
| Equation T1 | | $y = \left\vert A \cdot \left( 1 - 2B \cdot \exp\left(-\frac{TI}{T1}\right) \right) \right\vert $ | $y = \left\vert A \cdot \left( 1 - 2B \cdot \exp\left(-\frac{TI}{T1}\right) \right) \right\vert $ | $y = \left\vert A - B \cdot \exp\left(-\frac{TI}{T1}\right) \right\vert $ |
| Equation T2 | | $y = \left\vert A \cdot \exp\left(-\frac{TE}{T2}\right) \right\vert $ | $y = \left\vert A \cdot \exp\left(-\frac{TE}{T2}\right) \right\vert $ | $y = \left\vert A \cdot \exp\left(-\frac{TE}{T2}\right) \right\vert $ |

| **Segmentation** |  | **ICube (sC4)** | **MIRCEN (sC3)** |
| --- | --- | --- | --- |
| Brain Masking |  | Matlab / spm12 |  |
| Bias field correction |  | N4 |  |
| Registration |  | Non-rigid (ANTS (Avants et al., 2010)) | Rigid + Block-Matching affine (Lebenberg et al., 2010) |
| Number of atlas |  | 11 | 12 |
| Probability rule |  | Majority voting |  |

### MRI protocol

Acquisitions were performed on 7T horizontal scanners using the same MR sequences and parameters at centers C1 and C2 (aC1 and aC2 respectively, see details in Table 1). Preliminary *in vitro* experiments were performed at C1 and C2 in order to select the best sequences to use, with the objective to minimize acquisition time and geometric artifacts, and to maximize spatial resolution. A 3D MDEFT sequence (with Inversion Preparation as MPRAGE) was chosen for T1 mapping. Multi-Slice Multi-Echo (MSME) was chosen for T2 mapping. For T1 mapping, the MPRAGE sequence was run seven times with incremental inversion times (TI) (247, 408, 674, 1112, 1838, 3030, and 5000 ms). Repetition time (TR) was set to 6500 ms. For T2 mapping, a 3D MSME sequence with 28 echo times (TE), ranging from 8 to 224 ms, was used (DiFrancesco et al., 2008). TR was set to 600 ms. For both sequences, additional main parameters were: FOV: 2.7×2.7×2.8 cm^3^, matrix size: 128×128×66, spatial resolution: 211×211×424 μm^3^. Total experiment duration per animal was about 2 hours.

### Data analysis

#### Image processing workflow

Figure 1 illustrates the complete image processing workflow. Several preprocessing steps were performed using SPM12^1^ with MATLAB R2015a. Briefly, Bruker files were first converted into NIFTI images using home-made software. Three fitting procedures (fC1, fC2 and fC3) were used to obtain individual quantitative T1 and T2 maps. All anatomical images (T1-weighted with TI=247 ms) were rigidly realigned on a home-made rat template. Tissue segmentation was performed for each animal (Ashburner and Friston, 2005) based on the home-made tissue prior template. Using the Dartel registration algorithm (Ashburner, 2007), adapted for rat images, these tissue images were non-rigidly registered to create an average template.

**Figure 1:**
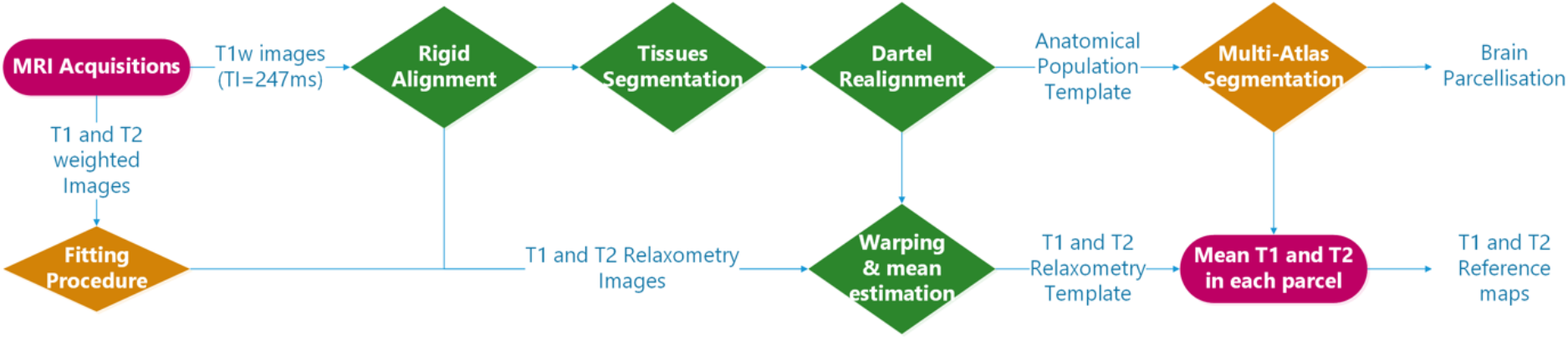
Processing workflow: the processing steps performed using SPM12 are shown in green. Three different fitting procedures and two implementations of the multi-atlas segmentation method were introduced.

The individual deformation field was then applied to the corresponding individual anatomical and relaxometry images, and all these images were averaged out to compute anatomical and relaxometry mean templates. The anatomical template was segmented using two multi-atlas approaches (sC3 and sC4) following a similar scheme as depicted in (Lancelot et al., 2014). Statistical analysis was performed with MS Excel 2010 and Real Statistics^2^. Because most of the samples did not present a normal distribution (Shapiro-Wilk test), non-parametric tests were performed. The fitting and multi-atlas segmentation algorithms were implemented within two platforms, namely BrainVISA^3^ for C3 and VIP^4^ (Glatard et al., 2013) for the other centers.

#### Fitting procedure

T1 and T2 weighted raw images were separately processed using three different fitting algorithms. The differences between these algorithms developed at C1, C2 and C3, respectively fC1, fC2 and fC3, are summarized in Table 1. Neither pre- nor post-processing were applied. All algorithms performed non-linear pixel-per-pixel fitting for each voxel independently. Negative values and values greater than 3000 ms for T1 and 300 ms for T2 were discarded. The optimization algorithm was Levenberg-Marquardt for fC1 and fC3 and the Simplex algorithm provided with ImageJ for fC2 (Nelder and Mead, 1965). All T2 estimation procedures rely on the same model equations.

#### Multi-Atlas Segmentation

Rat brain parcellation was performed using two multi-atlas approaches similar as the one proposed by Lancelot et al. (Lancelot et al., 2014), for a precise and reproducible delineation of brain structures in preclinical *in vivo* imaging. A maximum probability automatic delineation was obtained by the fusion of several manually delineated images placed in a common space and constituting the multi-atlas dataset. This dataset was registered to the native space of the MR image to segment. At each voxel, the most likely label in the dataset was selected by a maximum probability rule. Two versions of this approach were implemented, one at C3 (sC3) using BrainVISA and one at C4 (sC4) using VIP, which differ in some aspects as detailed in Table 1. Twenty-nine brain regions were defined (see Figure 2). MR data initially stored in SAS database were automatically sent to the VIP processing platform and processed results were seamlessly stored (see more details and the Figure 8 in (Commowick et al., 2018) for database and computing platform integration with Shanoir).

**Figure 2:**
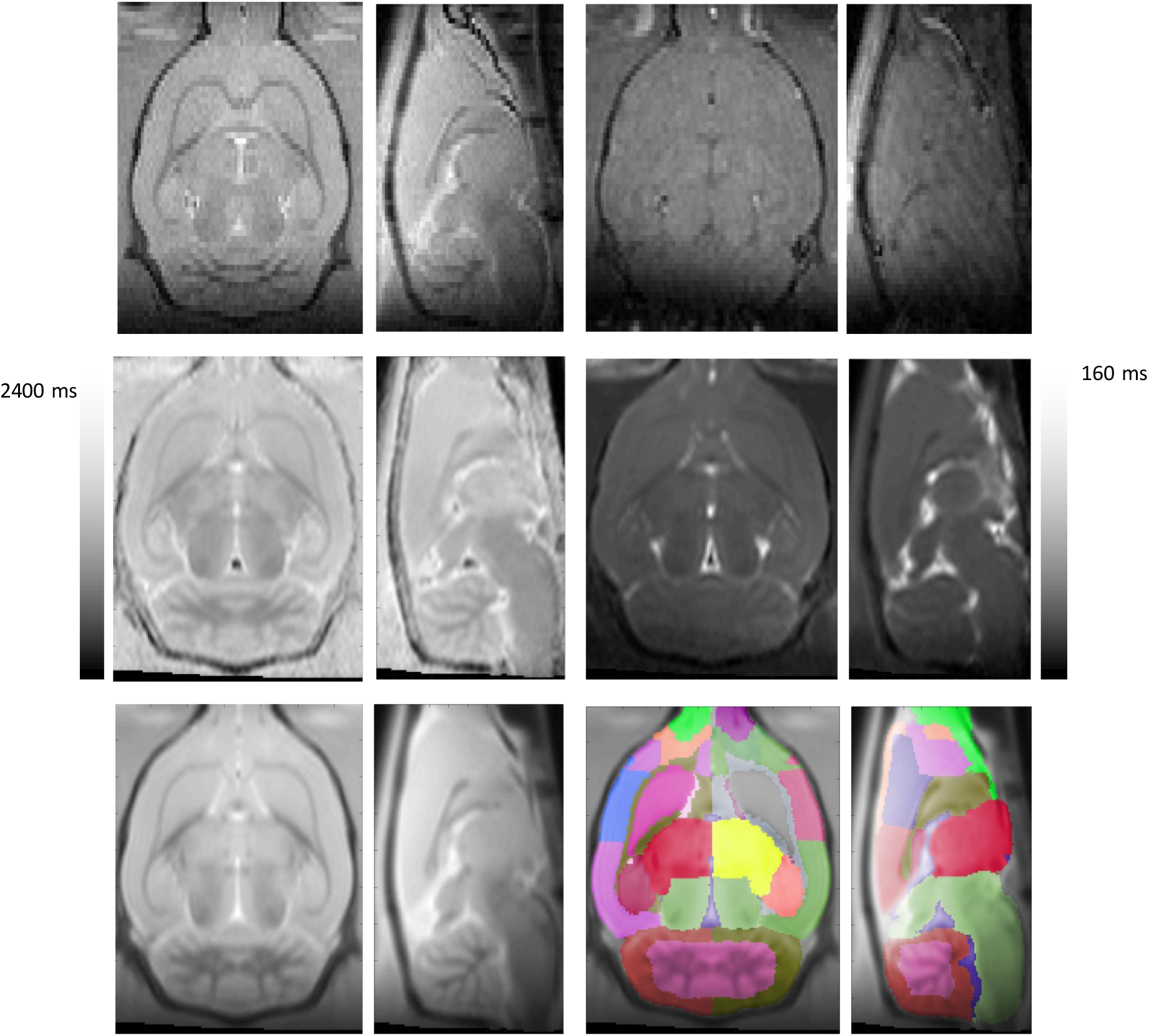
MR images processed at different steps of the processing pipeline. **Top**: Left. Individual raw T1-weighted sagittal and coronal images (TI: 247 ms). Right. Individual raw T2-weighted sagittal and coronal images (TE: 50 ms). **Middle**: Left. Sagittal and coronal views of the mean T1 relaxometry template (n = 40 rats). Right. Sagittal and coronal views of the mean T2 relaxometry template (n = 40 rats). **Bottom**. Left. Sagittal and coronal views of the T1-weighted template (40 rats; TI = 247 ms). Right. T1-weighted template parcellation (each brain region has a different color).

The twelve pipelines combining data (aC1 and aC2), fitting (fC1, fC2 and fC3) and segmentation (sC3 and sC4) were compared.

## RESULTS

### Processed Data

To illustrate the implemented processing pipelines, Figure 2 shows representative MR images of several processing steps.

### Inter-subject data variability

For each individual segmented rat brain, we computed the mean T1 and T2 relaxation times for the 29 regions (13 for each hemisphere and 3 unseparated). Figure 3 shows these values for each region of the left hemisphere of the 40 rats computed using the fC1 fitting pipeline and the sC4 multi-atlas segmentation.

**Figure 3:**
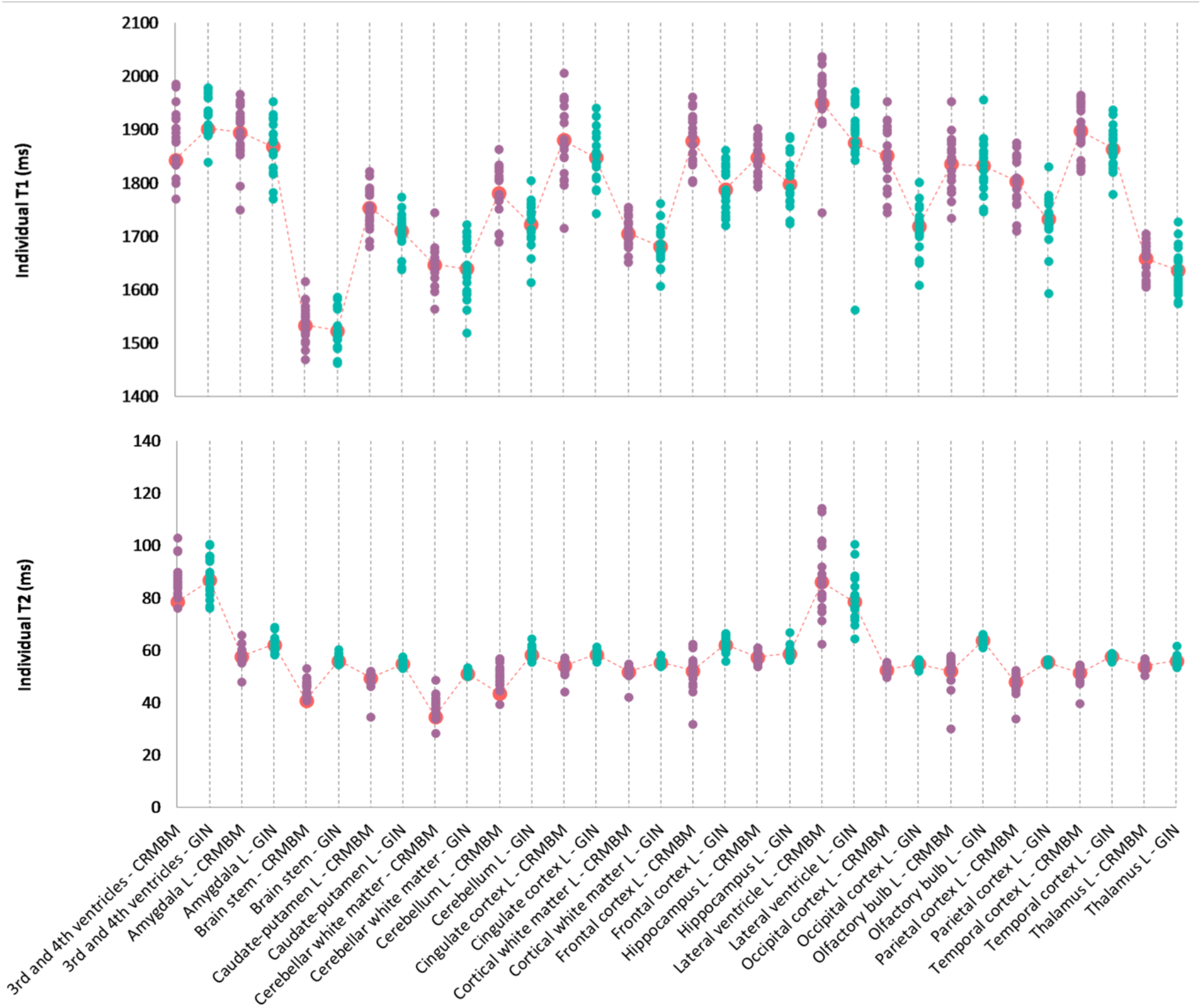
Individual relaxation time values for 16 regions of interest for the left hemisphere with 3 regions overlappping the two hemispheres. **Top:** Individual T1 values. **Bottom**: Individual T2 values. Blue circles for aC1 values; purple circles for aC2 values and corresponding mean values indicated with a red mark. fC1 fitting pipeline and sC4 multi-atlas segmentation were used.

We note that for T1 and T2 values, the dispersion is large for the ventricles (lateral, 3^rd^ and 4^th^ ventricles). On average, the differences between the minimum and maximum values of each region are 170 ms for T1 and 11 ms for T2 (left hemisphere excluding ventricles). We obtained similar results for the right hemisphere (169 and 9.3 ms respectively) and with the other pipelines (e.g. SM Figure 1 for fC2 fitting pipeline and sC3 multi-atlas).

### Inter-center acquisition variability

Figure 4 shows the differences in T1 and T2 values when computed using data acquired at aC1 vs at aC2 using the same pipeline.

**Figure 4:**
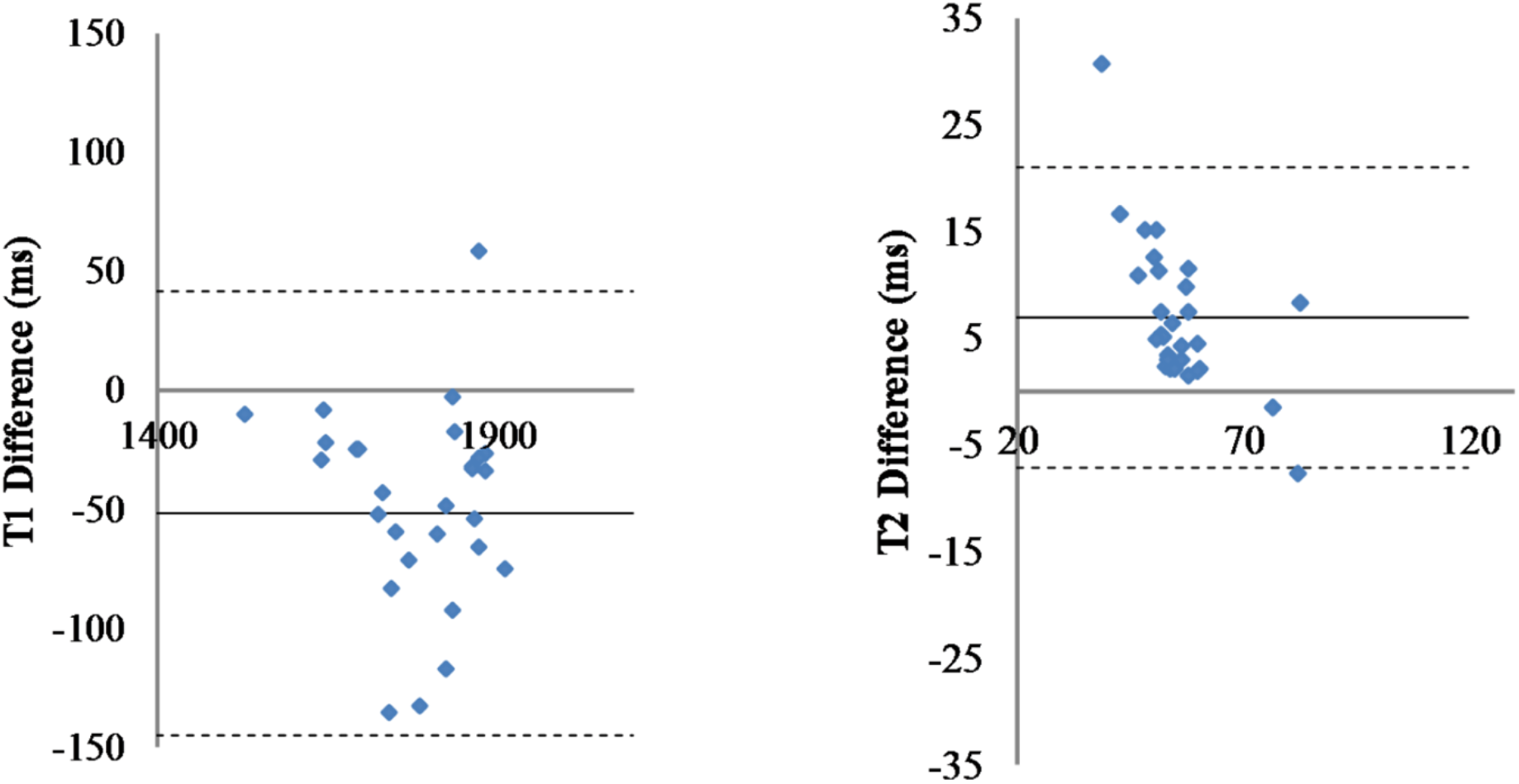
Inter-center variability. Differences for T1 (left) and T2 (right) relaxation time values computed from data acquired at aC1 (n=20) and aC2 (n=20) for the 29 regions of interest. Fitting pipeline fC1, Segmentation sC4. Solid line: mean difference. Dashed lines: ± 2 standard deviation. X-axis in ms.

The difference is significant between the T2 relaxation time values obtained from the two centers, but not for the T1 relaxation time values (Mann-Whitney test p=0.0002 with 9.52% mean error, and p=0.022 with 2.21% respectively). Similar results are obtained for the other fitting and segmentation method combinations (see SM Figure 2 for fC2 and sC3). Note that when fC3 fitting is used, differences between T1 relaxation time values become significant (see SM Figure 3).

### Intra-center acquisition reproducibility

Table 2 indicates the scan-rescan differences for each subject for T1 and T2.

**Table 2:** Mean scan-rescan differences.

|  | Subject 32 | Subject 21 | Subject 22 |
| --- | --- | --- | --- |
| **T1** | 22.6ms | 42.9ms | -25.8ms |
| **T2** | 2.59ms | -0.5ms | 0.05ms |

Figure 5 shows the differences for each subject in all ROIs of all subjects. Note that for T1, differences outside two standard deviations were found in the left lateral ventricle (Subject 32), the right olfactory bulb (Subject 21) and 3^rd^ and 4^th^ ventricles (Subject 22). For T2 (see Figure 6), similarly large differences were found in the left lateral ventricle (Subject 32), and the 3^rd^ and 4^th^ ventricles (Subject 21, Subject 22).

**Figure 5:**
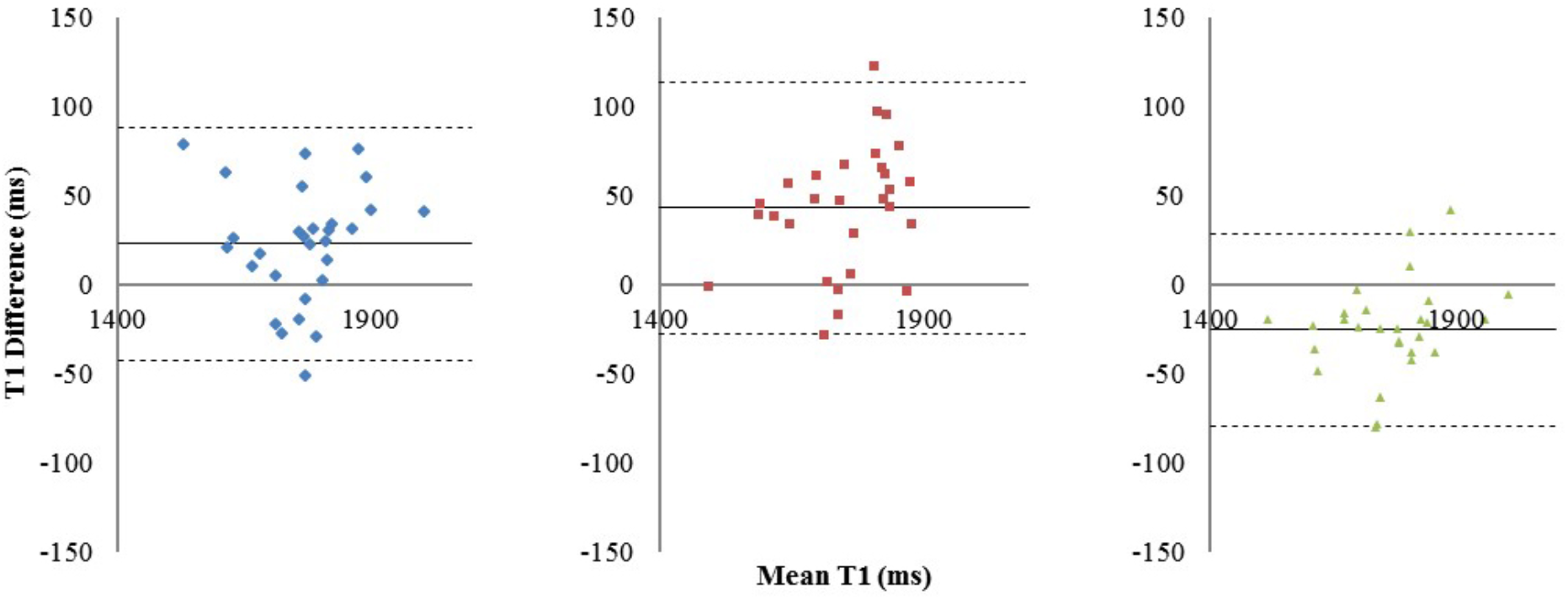
Intra-center reproducibility of T1 acquisition: Bland-Altman graph showing the differences in T1 relaxation time (mean values for each ROI) between the two acquisitions versus the corresponding mean T1 value. Left: Subject 32 (aC2). Middle: Subject 21 (aC1). Right: Subject 22 (aC1). Solid line: mean value, dashed lines ± 2 standard deviations. Fitting pipeline fC1, Segmentation sC4.

**Figure 6:**
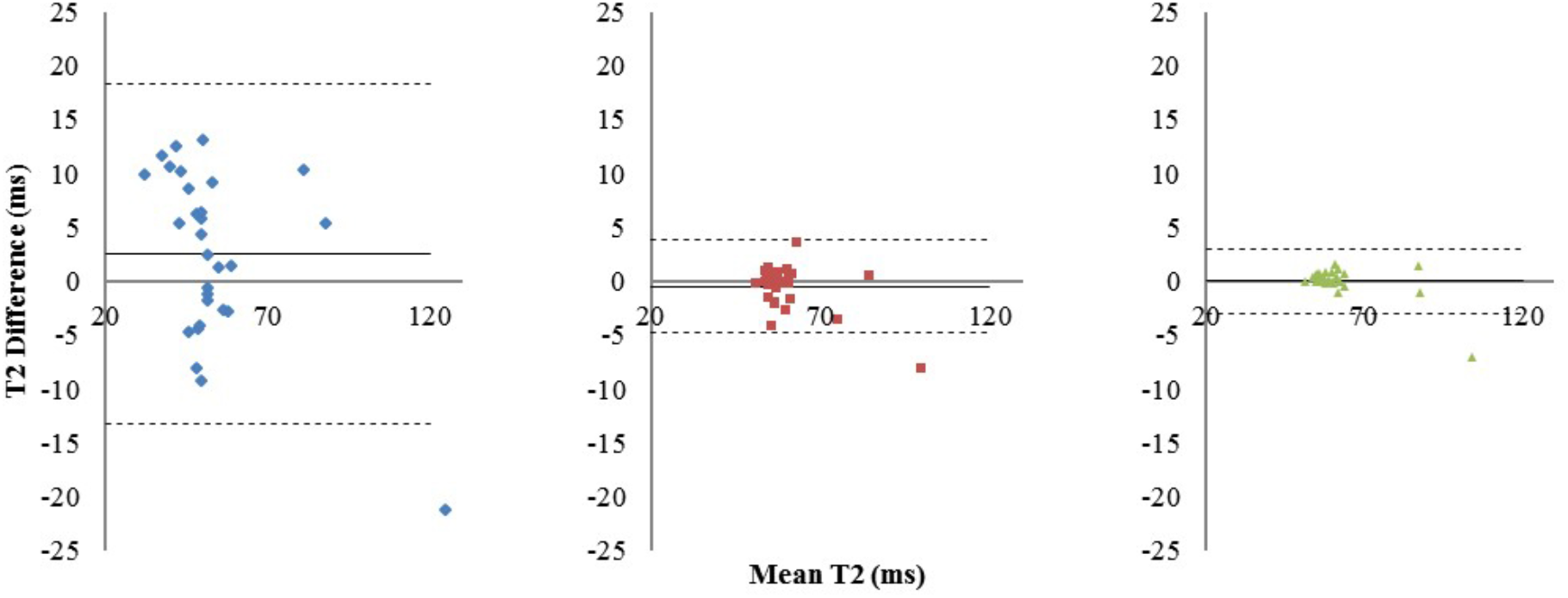
Intra-center reproducibility for T2: Bland-Altman graph showing the differences in T2 relaxation time (mean values for each ROI) between the two acquisitions versus the corresponding mean T2 value. Left: Subject 32 (aC2). Middle: Subject 21 (aC1). Right: Subject 22 (aC1). Solid line: mean value, dashed lines ± 2 standard deviations. Fitting pipeline fC1, Segmentation sC4.

Table 3 shows the parameters of the linear regression between the results from the scan-rescan experiment for each subject.

**Table 3:** Parameters for the linear regression in the scan-rescan experiment.

|  |  | Subject 32 | Subject 21 | Subject 22 |
| --- | --- | --- | --- | --- |
| T1 | Linear regression | $y = 0.930x$ | $y = 0,843x$ | $y = 0.906x$ |
|  |  | + 101.57 | + 235.05 | + 190.05 |
|  | R<sup>2</sup>: Coefficient of determination | 0.895 | 0.891 | 0,940 |
| T2 | Linear regression | $y = 1.154x$ | $y = 1.104x -$ | $y = 1.084x -$ |
|  |  | - 11.03 | 5.74 | 5.200 |
|  | R<sup>2</sup>: Coefficient of determination | 0.858 | 0.969 | 0,9905 |

Wilcoxon statistical test was run to compare the results from the first and second MR acquisition. P-values for T1 and T2 for each rat are summarized in Table 4.

**Table 4:** Wilcoxon test, p values.

|  | Subject 32 | Subject 21 | Subject 22 |
| --- | --- | --- | --- |
| **T1 map** | 0.479 | 0.089 | 0.266 |
| **T2 map** | 0.308 | 0.834 | 0.750 |

No differences are statistically significant. Same results were found with the other pipeline combinations (see SM Figures 4 and 5 and Table 2 for fC3 and sC3).

### Fitting pipeline comparison

To compare results obtained with the three fitting pipelines, regressions were computed and indicated a good consistency (see Figure 7). The linear regression parameters were y=0.9953x+5.11 (R^2^=0.9927) for T1 (C2f) vs T1 (C1f), y=0.9243x+111.9 (R^2^=0,9835) for T1 (C3f) vs T1 (C1f) and y=0.9315x+101.94 (R^2^=0.9969) for T1 (C3f) vs T1 (C2f).

**Figure 7:**
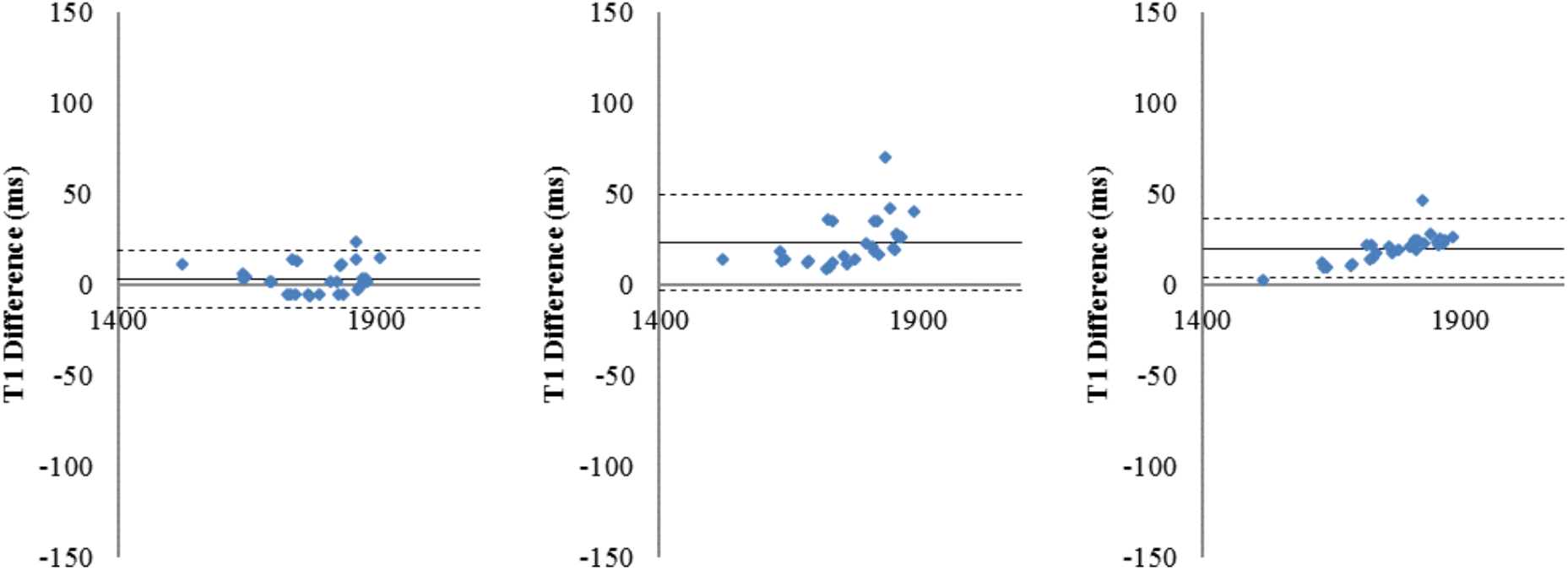
Comparison of fitting pipelines. T1 relaxation time differences found between different fitting pipelines for all regions of interest and averaged over the whole set of animals (n=40). Left: T1 differences for fC1 minus fC2; Middle: T1 differences for fC1 minus fC3; Right: T1 differences for fC2 minus fC3. sC4 for segmentation. Solid line: Mean difference. Dashed lines: ± two standard deviations.

**Figure 8:**
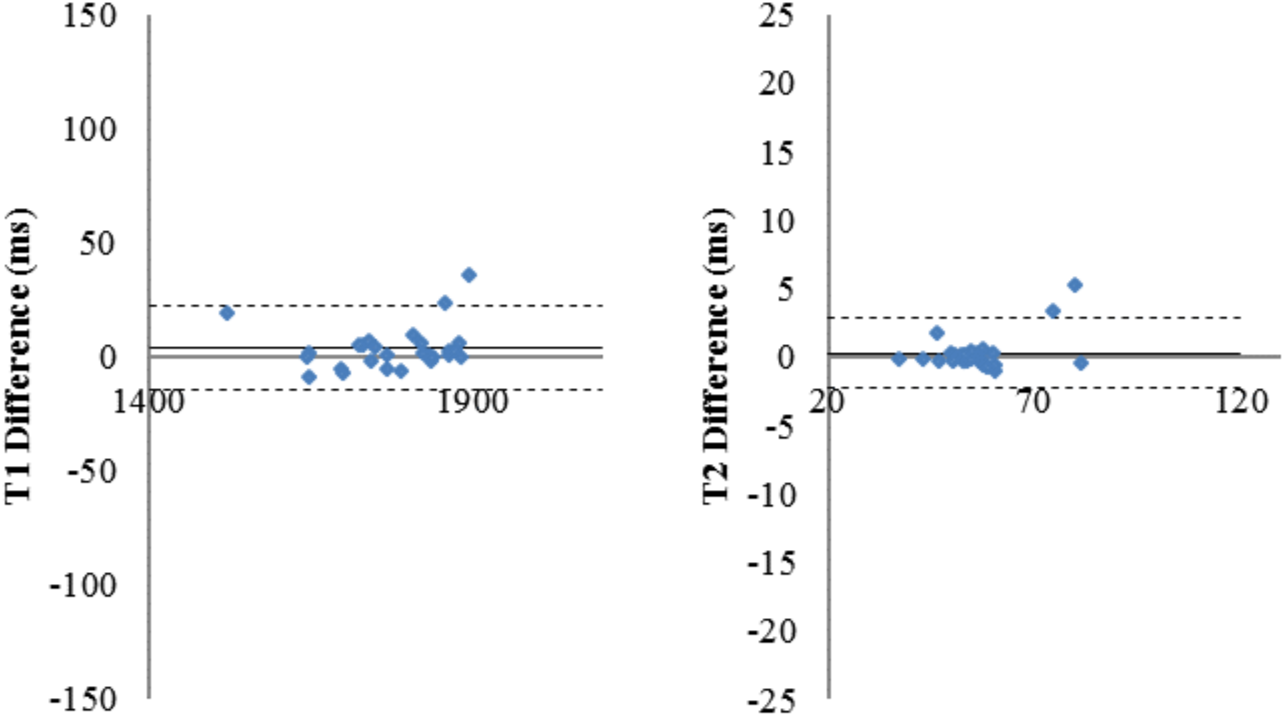
Comparison of segmentation pipelines. Bland-Altman graph of the T1 (left) and T2 (right) differences measured for all regions of interest and averaged over the entire set of animals (n=40) using the two different segmentation pipelines (one point per region of interest). fC1 fitting pipeline. Solid line: mean difference. Dashed line: ± two standard deviations.

Similar results were obtained for T2 relaxation times (see SM: Figure 6) and using sC3 (see SM: Table 3). Wilcoxon statistical test was run for each pair of pipelines. Table 5 reports the corresponding p-values (mean error to identity for T2: 3.33% for T1 1.04%).

**Table 5:** Wilcoxon test, p values, segmentation C4.

|  | C2 vs C1 | C1 vs C3 | C2 vs C3 |
| --- | --- | --- | --- |
| **T1** | 0,785 | 0,184 | 0,205 |
| **T2** | 0,858 | 0,273 | 0,380 |

### Segmentation pipeline comparison

To compare results obtained using the two segmentation pipelines, regressions were computed. As for the fitting pipelines, Figure 8 shows a good concordance between the methods. The linear regression parameters were y=1.012x-16.92 (R^2^=0.9902) for T1 (C3s) vs T1 (C4s) and y=1.0583x-2.9489 (R^2^=0.9865) for T2 (C3s) vs T2 (C4s).

P-values obtained using a Wilcoxon test were 0.823 and 0.994 for T1 and T2 relaxation times, respectively. These values were obtained via the two implementations of the same multi-atlas segmentation method (mean error to identity of 0.24% for T1and 0.81% for T2). Similar results were obtained when using the two other fitting methods (e.g. SM: Figure 7 for fC2).

### Comparison with the literature

Figure 9 shows the T1 and T2 values obtained in this study and those reported in the literature. In the case where literature only reported one cortical ROI, that value was replicated for all cortical ROIs of this study.

**Figure 9:**
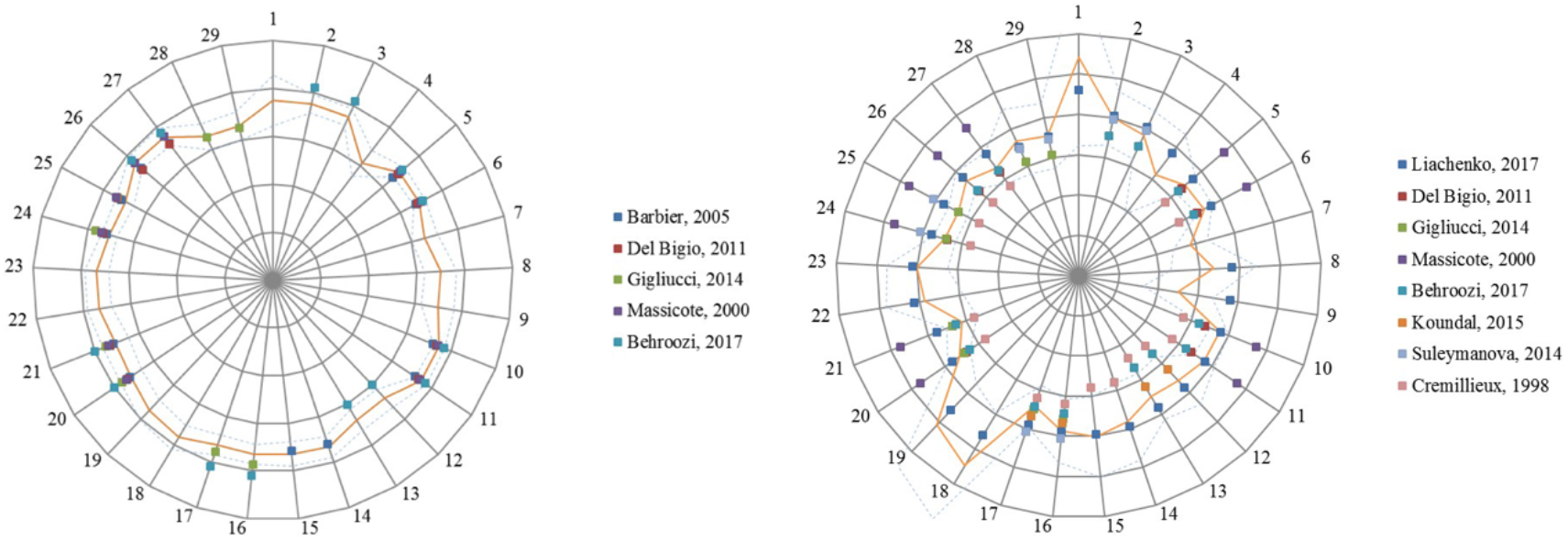
Relaxation values for this study and five published studies. (see References list for details) in 29 regions of interest. Left: T1 relaxation values. Scale: 500 ms / division. Right: T2 relaxation. Scale: 15 ms / division. Orange line indicates results from this study. Dashed lines indicate +/− one standard deviation from our results. 1: 3rd and 4th ventricles, 2: Amygdala L, 3: Amygdala R, 4: Brain stem, 5: Caudate-putamen L, 6: Caudate-putamen R, 7: Cerebellar white matter, 8: Cerebellum L, 9: Cerebellum R, 10: Cingulate cortex L, 11: Cingulate cortex R, 12: Cortical white matter L, 13: Cortical white matter R, 14: Frontal cortex L, 15: Frontal cortex R, 16: Hippocampus L, 17: Hippocampus R, 18: Lateral ventricle L, 19: Lateral ventricle R, 20: Occipital cortex L, 21: Occipital cortex R, 22: Olfactory bulb L, 23: Olfactory bulb R, 24: Parietal cortex L, 25: Parietal cortex R, 26: Temporal cortex L, 27: Temporal cortex R, 28: Thalamus L, 29: Thalamus R.

## DISCUSSION

Similarly to human population imaging, there are several well-founded motivations for animal population imaging: optimization of costs, reduction of experimentation duration, improvement of quality of science and enhancement of research discovery, notably by the use of sufficiently large animal cohorts for ensuring the validity of statistical results (see the special Lab Animal focus on reproducibility in animal research (Prescott and Lidster, 2017)). Although this domain is still in its infancy, we may expect that it will grow in the near future. Indeed, with the chosen use case, the multicenter computation of T1 and T2 relaxometry reference maps for rat brains at 7 Tesla, we clearly demonstrate the practicability and the potential of small animal population imaging.

The T1 and T2 relaxation times are tissue- and region-dependent, and because they may reflect micro-anatomical alterations, they constitute biomarkers for some pathologies (Deoni, 2010). To obtain such maps, weighted images are acquired with varying acquisition parameters such as echo time (TE), inversion time (TI) or flip angle. A voxel-per-voxel fitting, usually based on a three-parameter equation, provides T1 or T2 relaxation times (de Graaf et al., 2006, Guilfoyle et al., 2003, Wright et al., 2008). Interestingly, some new fingerprinting methods emerge based on dictionary use (Gao et al., 2015). Different monocentric studies were performed to measure T1 and T2 relaxation times in rodents, at different magnetic fields and using different acquisition protocols. They show that T1 relaxation time increases with magnetic field while T2 relaxation time decreases (de Graaf et al., 2006, Wright et al., 2008, van de Ven et al., 2007). Some values have been reported in the literature for few brain regions (Barbier et al., 2005, Behroozi et al., 2018, Cremillieux et al., 1998, Del Bigio et al., 2011, Gigliucci et al., 2014, Koundal et al., 2015, Liachenko and Ramu, 2017, Suleymanova et al., 2014), but no consensus has been reached yet about values of reference for specific rat brain regions. To define such reference maps, a large number of brain structures or regions should be considered and a sufficient number of animals should be included to reflect inter-individual variability. In this context, a multicenter study is relevant. In this study, four centers were involved, two of which also contributed acquired data to evaluate the feasibility of pooling data. The influence of image processing steps on the final maps was assessed by comparing three fitting algorithms provided by three centers. Finally, to derive relaxation times per brain areas, two multi-atlas segmentation pipelines from two centers were considered.

For Human studies, to facilitate data storage, data sharing and data processing based on specific customized pipelines, different infrastructures have been proposed such as for instance COINS (Landis et al., 2016), LORIS+BRAIN (Das et al., 2016) or LONI (Rex et al., 2003) for Neuroimaging multicenter studies (see recent works in this field (Dojat et al., 2017)). These infrastructures support the “Open Science” approach, an international action to improve the use of resources, to ease study replication and then to strengthen the validity of scientific results (NAP, 2018). This promotes studies on very large cohorts e.g. (Adhikari et al., 2018), the development of reference databases (see for Alzheimer disease (Li et al., 2017), Parkinson disease (Chahine et al., 2018) or Human connectome project (Hodge et al., 2016)) and fair and robust comparison of image processing solutions (Commowick et al., 2018). Here, we propose to use the extension of the SHANOIR environment (Barillot et al., 2016) for storing preclinical imaging data, coupled with the VIP architecture for the dedicated image processing pipelines execution (Glatard et al., 2013).

First, we explored the quality of data acquired on different animals at different centers. We evaluated the data reproducibility (intra- and inter-center differences) to assess whether data pooling was feasible.

### Acquisition reproducibility

Figures 5 and 6 show good intra-center reproducibility (n=3) for T1 and T2 values, with less dispersion for the latter at C1 (see Tables 2 and 3). There was no significant difference in the scan-rescan experiment of each rat. Some regions were more prone to measurement errors, especially the ventricles; certainly, due to small realignment errors and large intensity variations between ventricles and adjacent ROIs (see below). Good reproducibility was also obtained when using different processing solutions (SM: Figures 4 and 5).

### Data pooling

We successfully combined MR data acquired on forty rats in two different centers, GIN (n=20) and CRMBM (n=20). As for multicenter studies in Humans, we strongly limited the inter-center data variability by adopting the same image acquisition conditions (same rat strain and same animal provider, same magnetic field, same receive coil, same sequence parameters and same operator). We took advantage of the fact that the fleet of MR scanners for preclinical studies tends to be more uniform across labs than Human imaging systems. Then, the MR scanners in both centers were designed by the same manufacturer (Bruker Biospin) and, as indicated in Table 1, there were very few differences between the two systems. This way, the acquisition sequence parameters could also be set identical in both centers. Consequently, Figure 3 shows a weak dispersion of individual T1 and T2 values in brain regions, except for the ventricles (lateral and 3^rd^ & 4^th^ ventricles). This could be a consequence of the small number of voxels of these structures compared to other regions (3500 and around 11000 voxels for lateral and 3^rd^ and 4^th^ ventricles, respectively, versus 33000 voxels on average for the other structures) and to the large differences in relaxation times between brain tissue and cerebrospinal fluid. Moreover, contrast in T1-weighted imaging between ventricles and tissue is low. This renders the registration process more prone to errors. Ventricles are filled with cerebrospinal liquid. During acquisition, small movements of cerebrospinal fluid may lead to a biased estimation of relaxation times in the ventricles (especially for T2 which exhibits the largest difference between tissue and ventricle). Moreover, for some rats (n=3), ventricles were dilated. Such dilatation may impact the global results (Massicotte et al., 2000). No significant differences were detected for T1 values when computed from data acquired at C1 or C2 for different fitting and segmentation pipelines (see Figure 4 and SM: Figure 2) but for C3 fitting algorithm (p<0.01, SM: Figure 3) T2 differences were significant for all conditions. We note that, in the literature, T2 measurements seem more variable between centers than T1 (see Figure 9). After examining multicenter data quality, we searched for possible differences when processing data with software developed in different centers.

### Image processing pipeline comparison

The goal was not to compare the processing solutions between centers for selecting the best one. We rather explored whether data processing can be distributed in different centers using the solution locally available. For relaxometry map computation, two main steps are required: fitting the raw data to extract relaxation times and segment the brain to compute mean values per brain area. Fitting data is clearly a simple mathematical operation. No consensus exists for this operation to map relaxometry and, as indicated in Table 2, differences exist in available solutions hosted in our three centers. Figure 7 shows that the differences in the fitting equation and optimization methods used did not impact the final results. Note that differences generated by the use of different fitting parameters were lower than differences generated by different data acquisition centers (compare Figure 7 with Figure 4) and differences between data acquired at the same center (compare Figure 7 with Figure 5). This is similar for T2 values (see SM Figure 6). For brain segmentation, we adopted a multi-atlas procedure. Multi-atlas techniques outperform single atlas approaches for accounting individual structural variability (Wang et al., 2014). Here, we considered two implementations on two different executing platforms, BrainVISA and VIP, of the multi-atlas approach proposed in (Lancelot et al., 2014) for rats. As mentioned in Table 1, some preprocessing steps differ: registration (ANTS (Avants et al., 2010) vs block-matching combined to Free Form Deformation (Lebenberg et al., 2010)) and the number of atlas used (11 vs 12). Comparison of the parcellation obtained using the two methods revealed no differences in term of volume for each parcel and DICE score which were higher than 0.8 but for ventricles (see SM: Figure 8). As indicated in Figure 8 (and SM: Figure 7), no differences were significant between T1 and T2 relaxation values obtained using the two multi-atlas segmentation procedures.

### Reference Maps

Publicly available quantitative reference maps for longitudinal (T1) and transverse (T2) relaxation times are valuable to investigate and monitor deviations from normality in animal model of pathologies. Some T1 and T2 relaxation values have been reported in the literature based on a limited number of animals. We extended these works by using more animals (40) and exploring additional regions. Figure 9 shows the T1 and T2 relaxation values obtained in this study together with literature values. There is a good consistency between all T1 relaxation values. This is consistent with the low dispersion we measured between two centers (2%). The dispersion is larger for T2 relaxation values especially for older studies. We reported a 9.5% dispersion between our two data provider centers. Altogether, reported values are in good agreement with several recent studies (Del Bigio et al., 2011, Gigliucci et al., 2014, Koundal et al., 2015, Liachenko and Ramu, 2017, Suleymanova et al., 2014).

### Some limitations

Data were acquired in only one animal strain (Sprague Dawley) and at one age (young adult). These two factors may modify relaxation times (https://www.ncbi.nlm.nih.gov/pubmed/29060536, https://www.ncbi.nlm.nih.gov/pubmed/27379992, https://www.ncbi.nlm.nih.gov/pubmed/30302499, https://www.ncbi.nlm.nih.gov/pubmed/27333937). Finer changes may also occur in case of physiological changes (perfusion, oxygenation). In addition, our reference maps could be refined by including more individuals.

In conclusion, we have demonstrated the feasibility of multicenter preclinical studies, sharing raw data and processing pipelines between distributed centers. Similar relaxometry maps for different rat strains and at different ages would also be relevant and could be added to the data obtained in this study. Eventually, differences between normal and pathological models could be explored. Raw data and reference T1 and T2 relaxometry maps are therefore freely available on request (https://shanoir.irisa.fr/Shanoir/login.seam, as well as processing pipelines hosted here https://www.creatis.insa-lyon.fr/vip/, contact M. Dojat). Please refer to the present paper in case of the reuse of these datasets and pipelines.

## Acknowledgements

The authors thank the technical staff of VIP (https://www.creatis.insa-lyon.fr/vip/), for their support for the multi-atlas segmentation algorithm portage, and the FLI-IAM team (https://portal.fli-iam.irisa.fr/preclinical_imaging_studies) for data management support.

## Footnotes

1 http://www.fil.ion.ucl.ac.uk/spm

2 Real Statistics Resource Pack software (Release 5.4). Copyright (2013 – 2018) Charles Zaiontz. www.real-statistics.com

3 http://brainvisa.info/

4 Virtual Imaging Platform https://www.creatis.insa-lyon.fr/vip/

