## Supplementary material for "A Multicenter Preclinical MRI Study: Definition of Rat Brain Relaxometry Reference Maps": SM

### Table 1: Animal information

| Acq month | center | Rat_id | Weight (g) | Breath rate (bpm) | Temperature (°C) |
| --- | --- | --- | --- | --- | --- |
| December | GIN | S1 | 274 |  |  |
| December | GIN | S2 | 258 |  |  |
| December | GIN | S3 | 286 |  |  |
| December | GIN | S4 | 264 |  |  |
| December | GIN | S5 | 300 |  |  |
| December | GIN | S6 | 294 | 70-82 |  |
| December | GIN | S7 | 304 | 75-97 |  |
| December | GIN | S8 | 304 | 82 |  |
| December | GIN | S9 | 288 | 70-80 |  |
| December | GIN | S10 | 296 | 75-97 |  |
| December | CRMBM | S11 | 249,5 | 60 |  |
| December | CRMBM | S12 | 263,6 | 80-88 | 36,8 |
| December | CRMBM | S13 | 267 | 30 | 34 |
| December | CRMBM | S14 | 269 | 75 | 37,2 |
| December | CRMBM | S15 | 273 | 65-75 | 37 |
| December | CRMBM | S16 | 279,9 | 90 | 37,5 |
| December | CRMBM | S17 | 277 | 80 | 37,7 |
| December | CRMBM | S18 | 277 | 75-80 | 35,6 |
| December | CRMBM | S19 | 271,3 | 65 | 34,6 |
| December | CRMBM | S20 | 274 | 70 | 35,9 |
| May | GIN | S21 | 294 | 70 | 37,7 |
| May | GIN | S22 | 256 | 70-75 | 37,8 |
| May | GIN | S23 | 278 | 70 | 38,5 |
| May | GIN | S24 | 282 | 70 | 37,9 |
| May | GIN | S25 | 268 | 65 | 36,8 |
| May | GIN | S26 | 274 | 70 | 37,7 |
| May | GIN | S27 | 314 | 70 | 37,5 |
| May | GIN | S28 |  | 75 | 37,1 |
| May | GIN | S29 | 262 | 60 | 36,8 |
| May | GIN | S30 | 274 | 65 | 37 |
| May | CRMBM | S31 | 297,5 | 60 | 34,8 |
| May | CRMBM | S32 | 284,9 | 60 | 34,8 |
| May | CRMBM | S33 | 302,7 | 55-55 | 35,3 |
| May | CRMBM | S34 | 284,8 | 55-60 | 35,1 |
| May | CRMBM | S35 | 290,5 | 70-60 | 35 |
| May | CRMBM | S36 | 284,3 | 65 | 34,4 |
| May | CRMBM | S37 | 275,7 | 60 | 34,5 |
| May | CRMBM | S38 | 286,9 | 120-60 | 34,8 |
| May | CRMBM | S39 | 285,5 | 120-60 | 36,2 |
| May | CRMBM | S40 | 290 | 65 | 34,3 |
| Mean |  |  | 279.40 (SD=13.87) | 67.13 (SD=11.42) | 36.21 (SD=1.36) |

#### Data

**Inter subject variability**

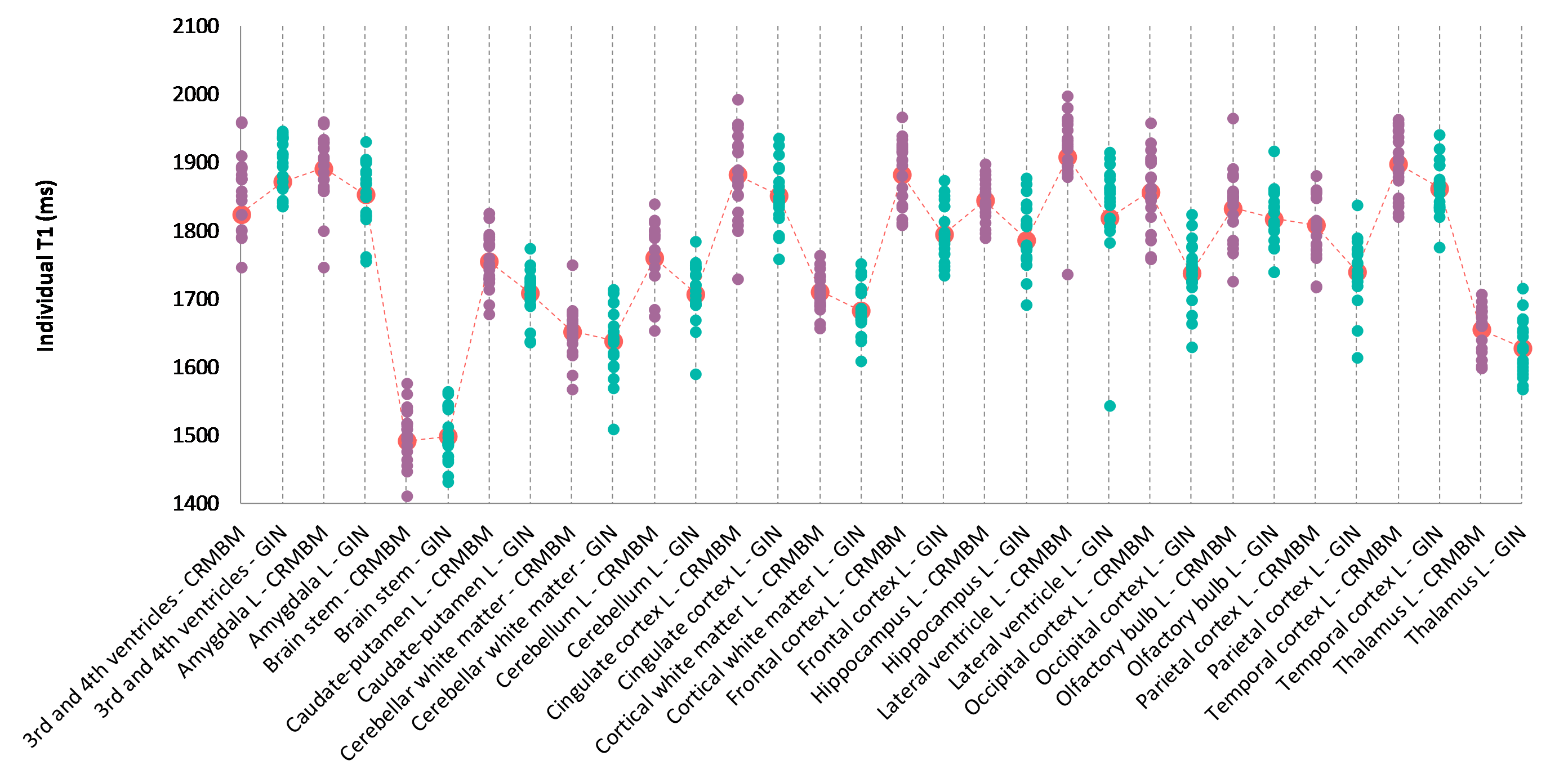

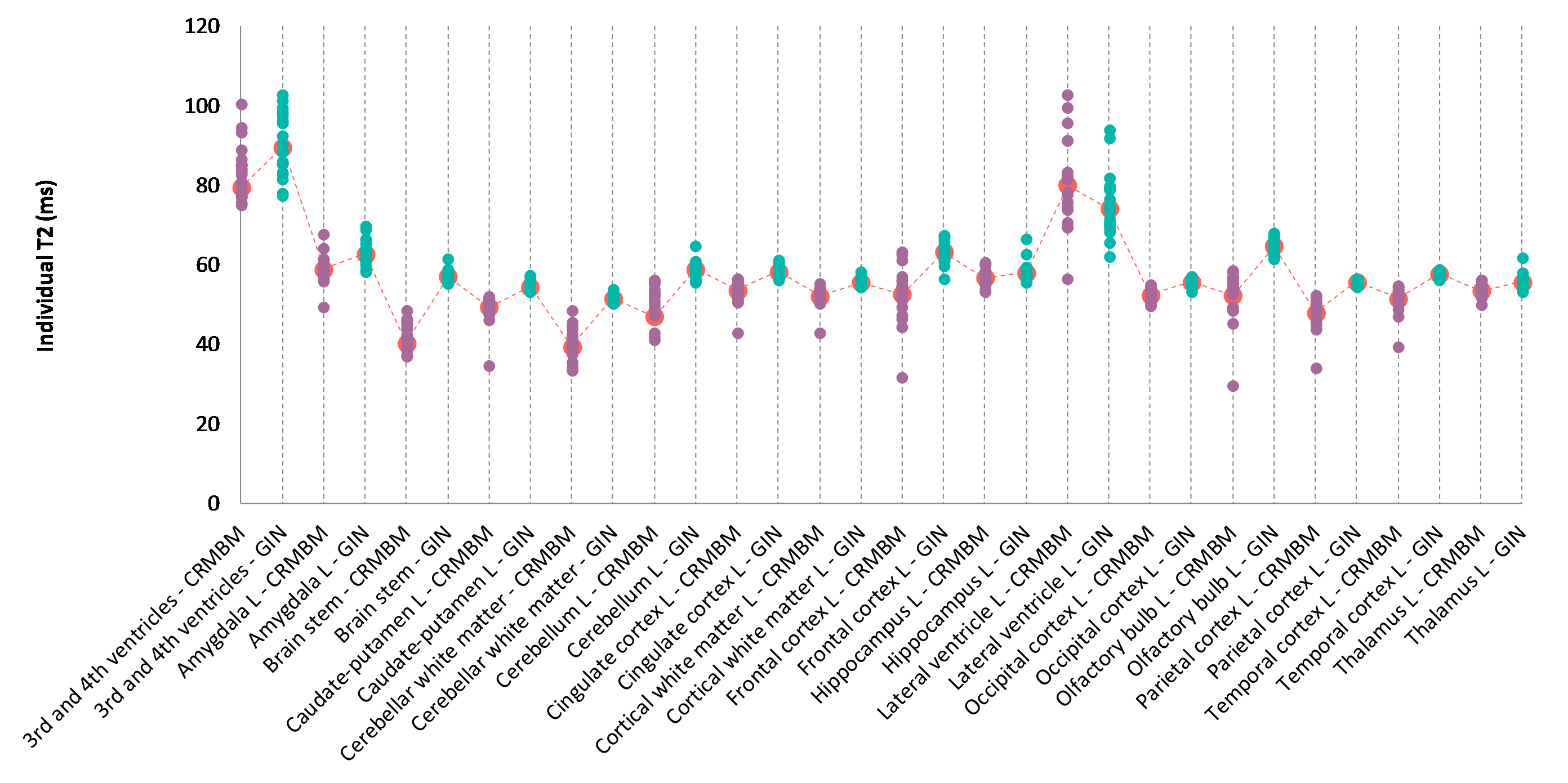

Figure 1: Individual relaxation time values for 13 regions of interest for the left hemisphere. Top: Individual T1 values. Bottom: Individual T2 values. Blue circles for aC1 values; purple circles for aC2 values and corresponding mean values indicated with a red mark. fC2 fitting pipeline and sC3 multi-atlas segmentation were used.

**Inter-center variability**

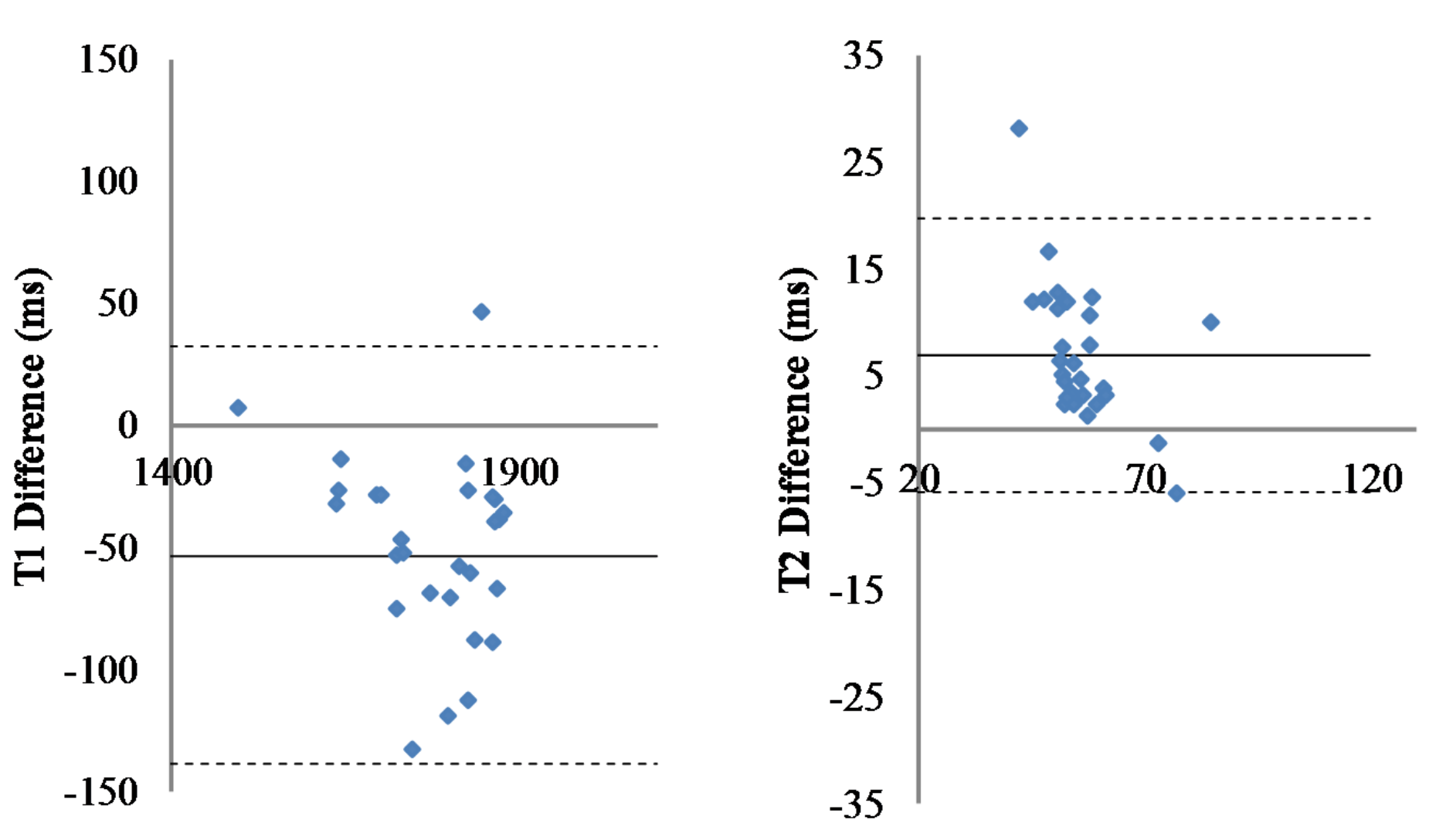

Figure 2: Inter-center variability. Differences for T1 (left) and T2 (right) relaxation time values computed from data acquired at C2 (n=20) and C1 (n=20) for the 29 regions of interest. Fitting pipeline fC2, Segmentation sC3. Solid line: Mean difference. Dashed lines: ± 2 standard deviation.

Mann-Whitney test **p=9,80 10-5** and p=0.010 with 9.40% and 2.26% mean error to identity for respectively T2 and T1 relaxometry time values.

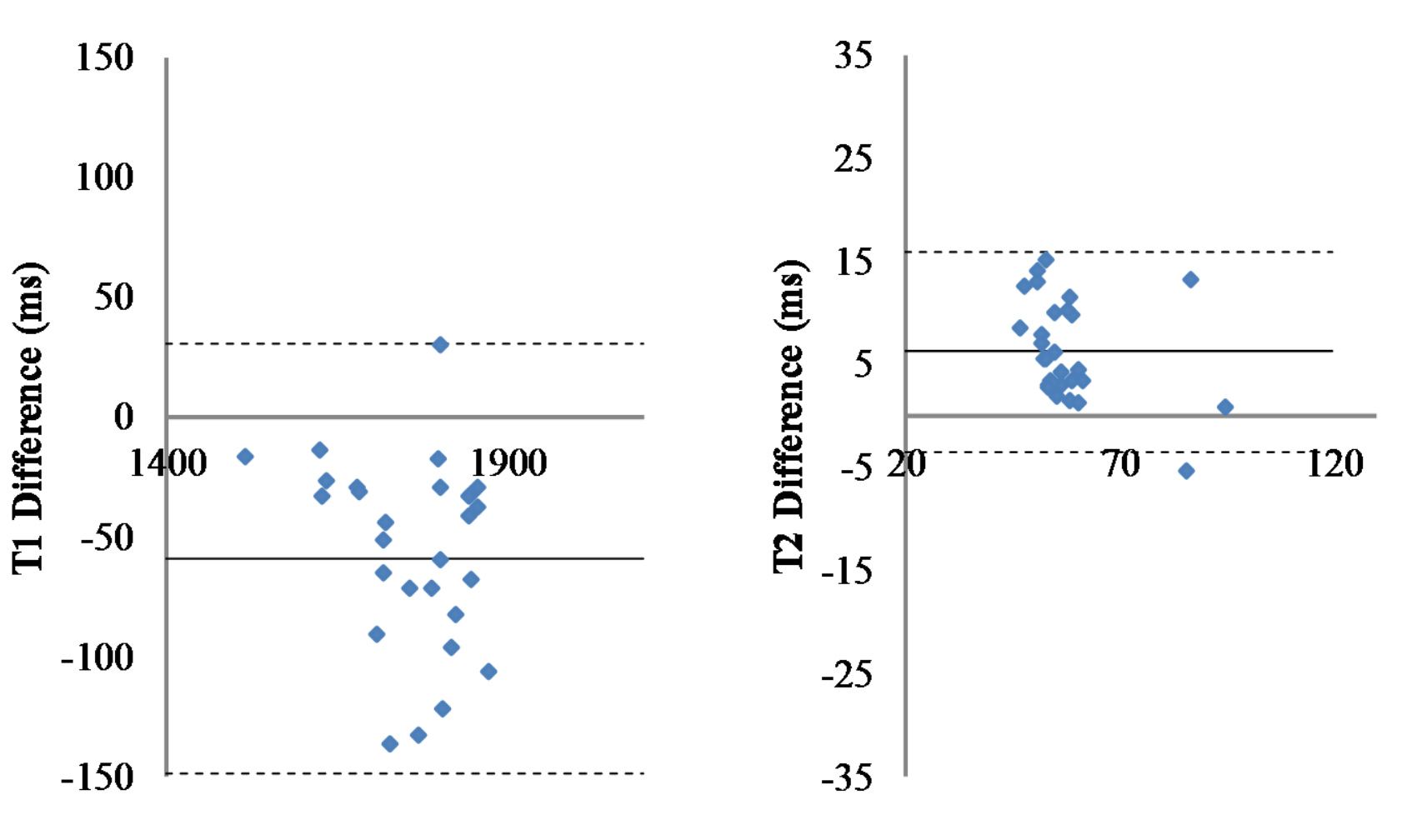

Figure 3: Inter-center variability. Differences for T1 (left) and T2 (right) relaxation time values computed from data acquired at C2 (n=20) and C1 (n=20) for the 29 regions of interest. Fitting pipeline fC3, Segmentation sC4. Solid line: Mean difference. Dashed lines: ± 2 standard deviation.

Mann-Whitney test **p=9,19 10-5** and **p=0.0063** with 8.07% and 2.46% mean error to identity for respectively T2 and T1 relaxometry time values.

#### Reproducibility

Table 2: Wilcoxon test. p values

|  | Subject32 | Subject21 | Subject22 |
| --- | --- | --- | --- |
| T1 map | 0,529 | 0,055 | 0,294 |
| T2 map | 0,379 | 0,773 | 0,613 |

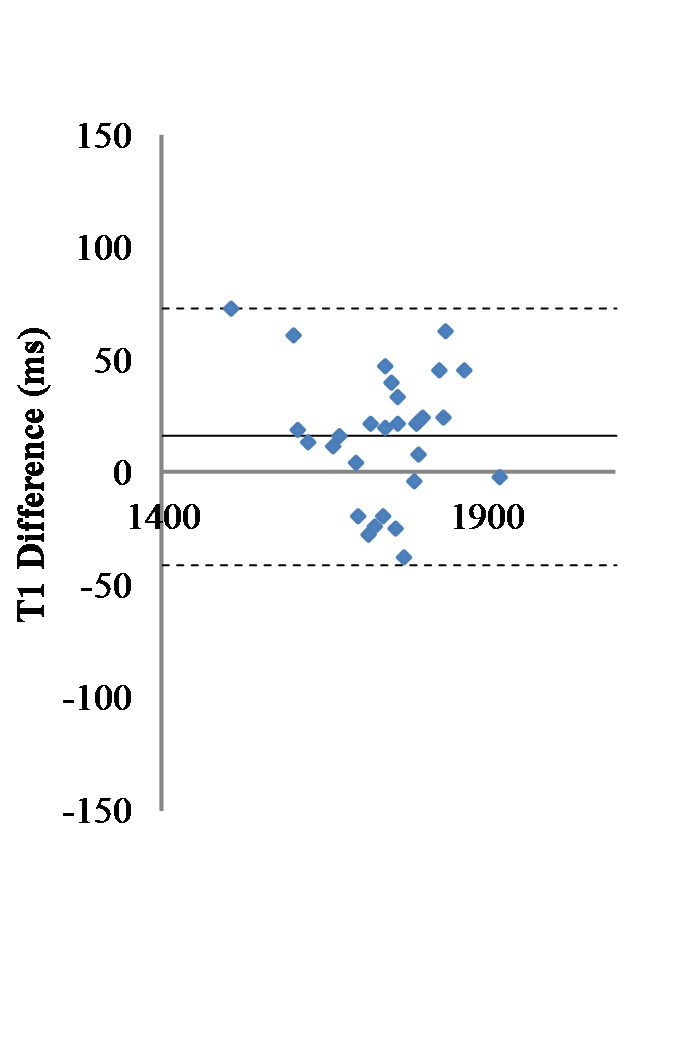

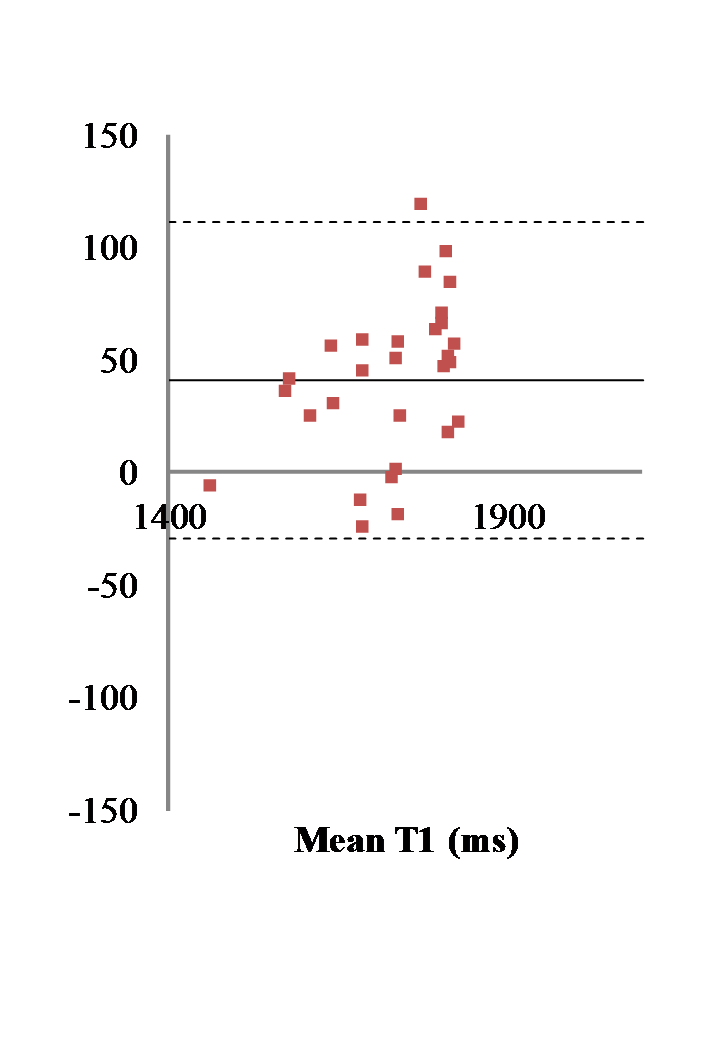

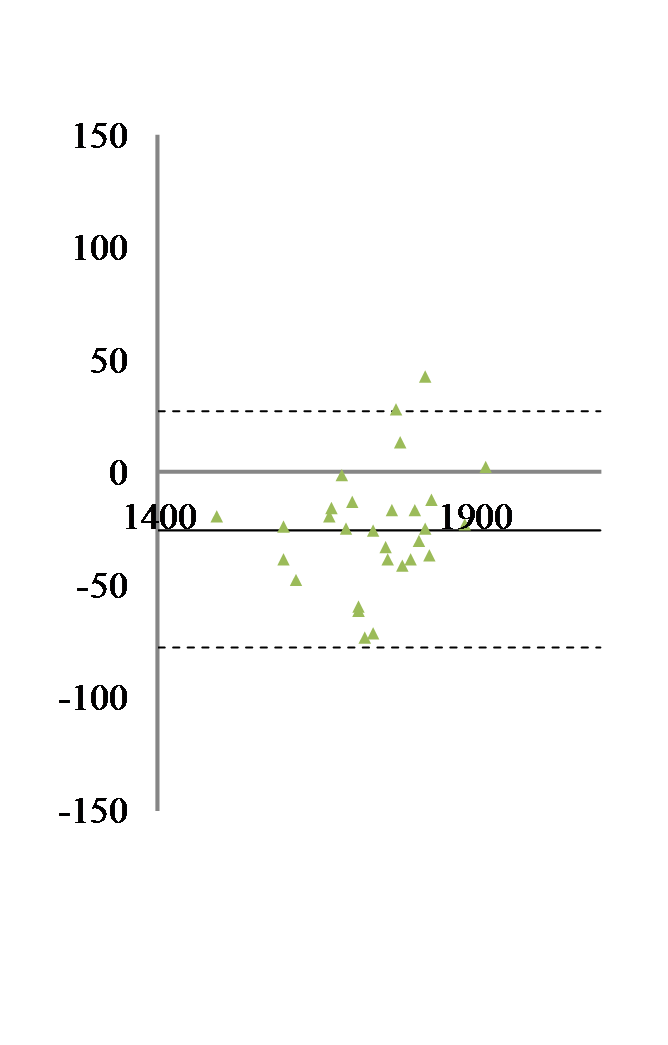

Figure 4: Intra-center reproducibility for T1: Differences for T1 values for each ROI between the two acquisitions versus the corresponding mean T1 value. Left: S32 (C2). Middle S21 (C1). Right S22 (C1). Bold line: mean value, dashed lines +/- 2 standard deviations. Fitting pipeline fC3, Segmentation sC3.

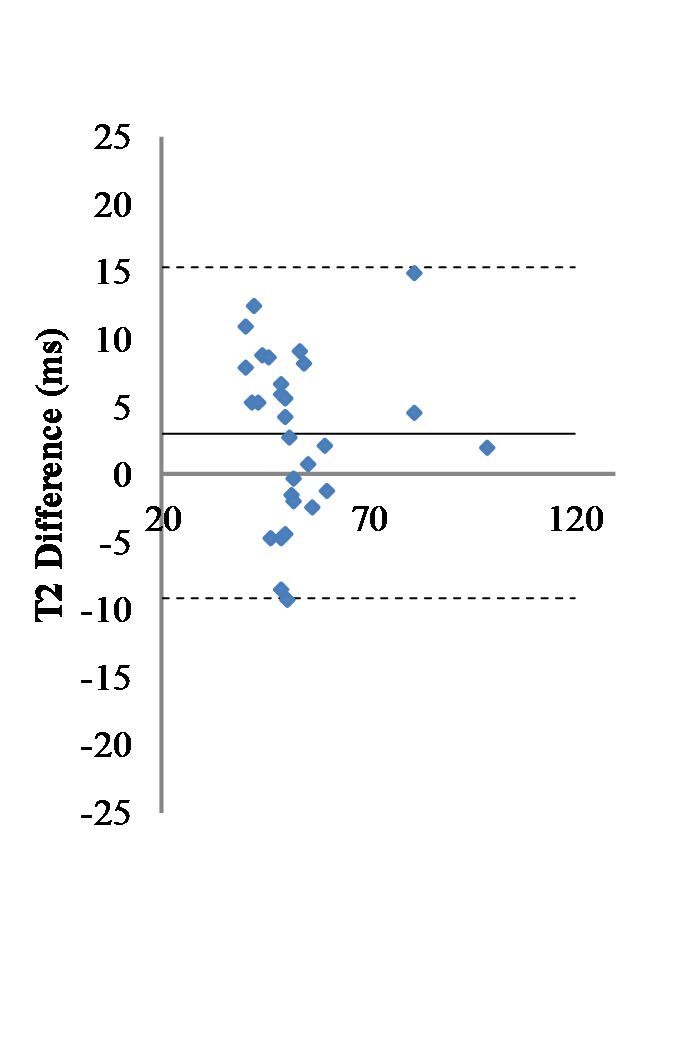

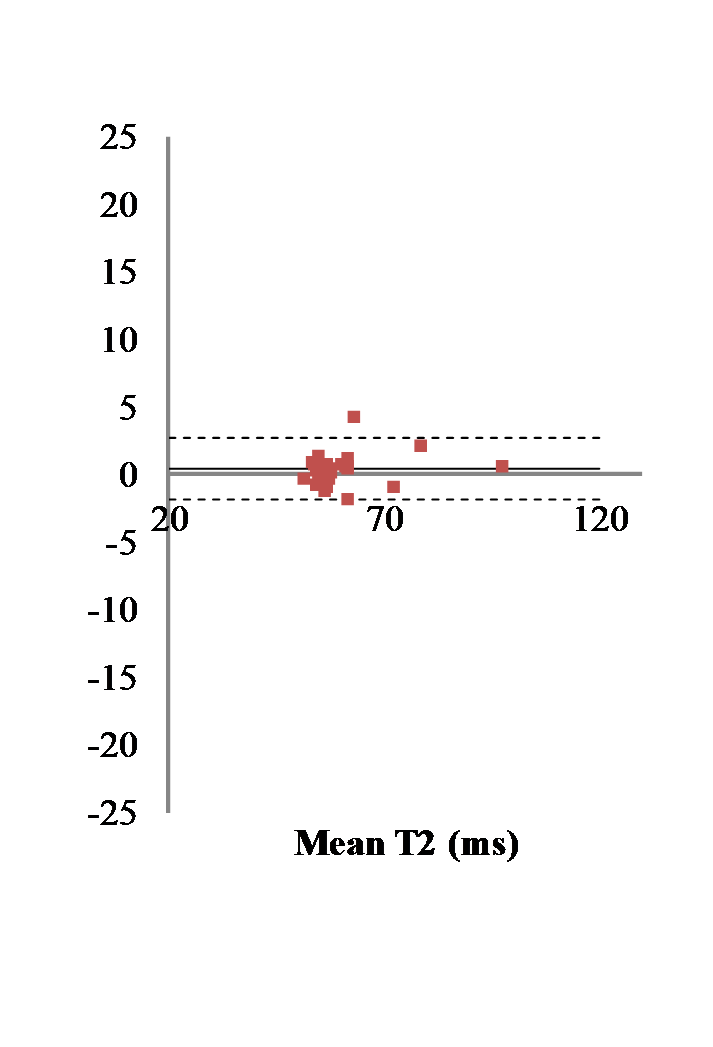

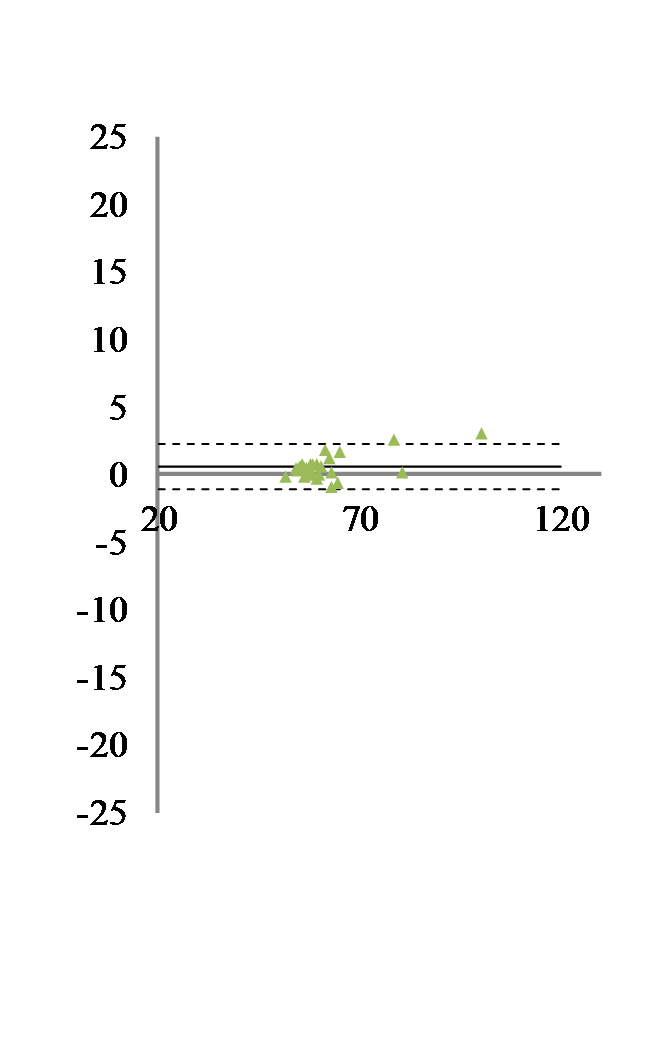

Figure 5: Intra-center reproducibility for T2: Differences for T2 values for each ROI between the two acquisitions versus the corresponding mean T2 value. Left: S32 (C2). Middle S21 (C1). Right S22 (C1). Bold line: mean value, dashed lines +/- 2 standard deviations. Fitting pipeline fC3, Segmentation sC3.

#### Fitting pipelines comparison

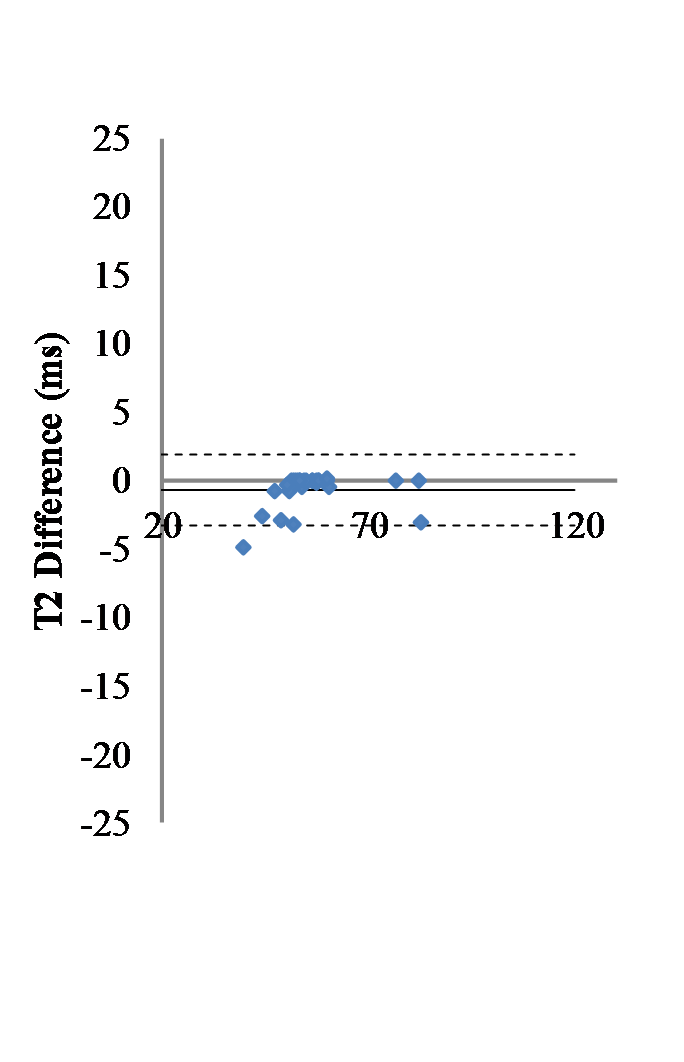

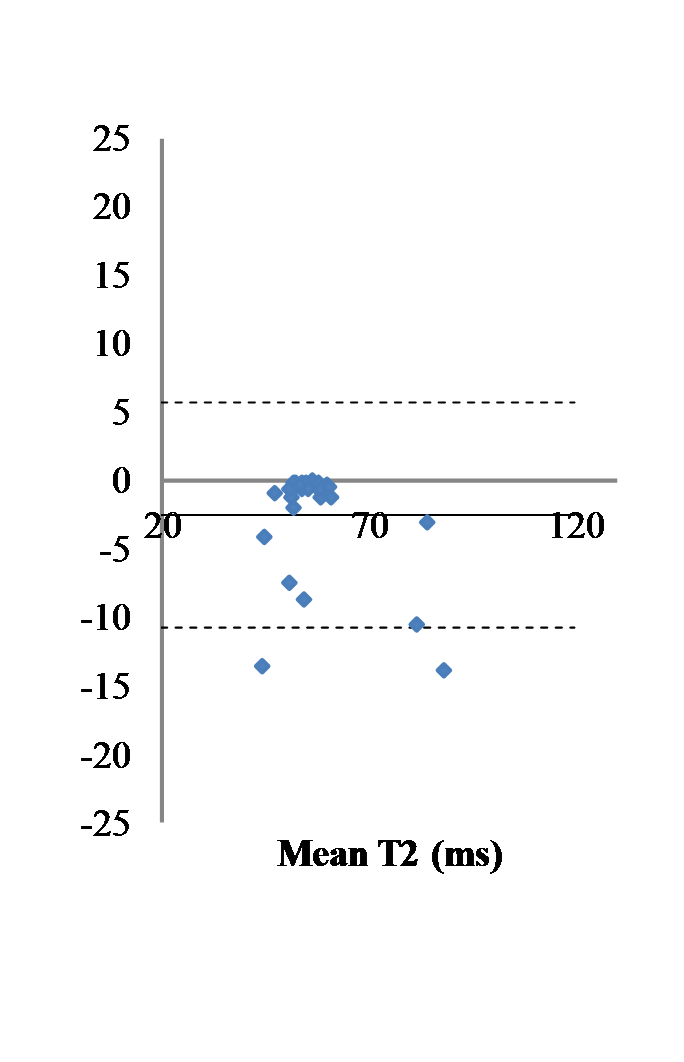

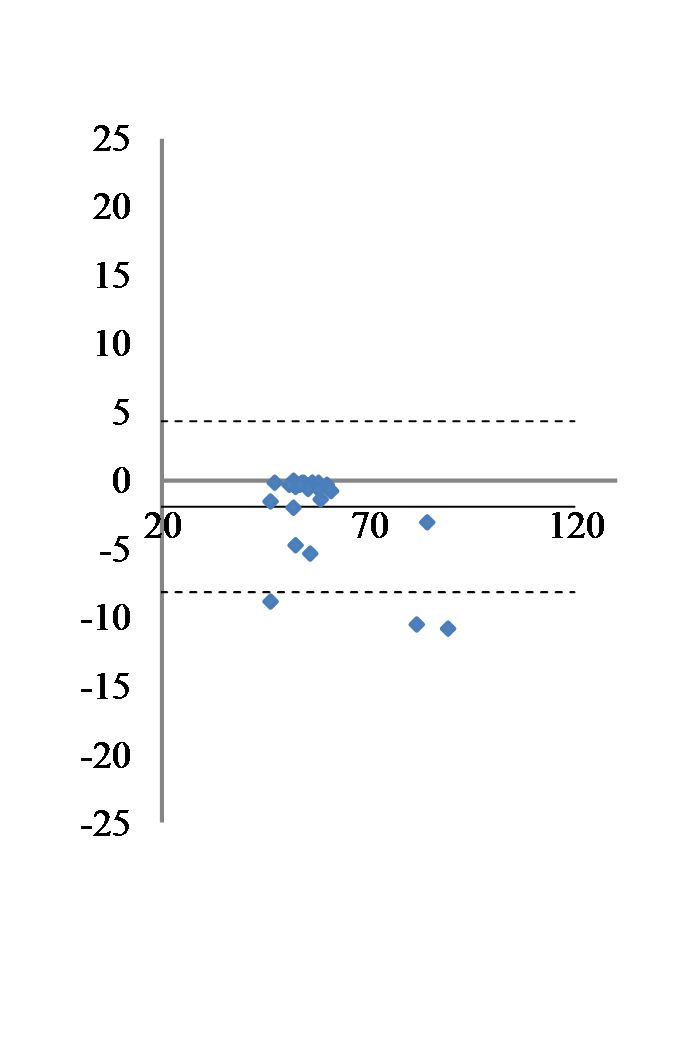

Figure 6: Comparison of fitting pipelines. T2 relaxation times values differences computed using different fitting pipelines for all regions of interest and average out over the whole set of animals (n=40). Left: T2 values differences for fC1 minus fC2; Middle: T2 values differences for fC1 minus fC3; Right: T2 values differences for fC2 minus fC3. sC4 for segmentation. Solid line: Mean difference. Dashed lines: ± two standard deviations.

Table 3: Wilcoxon test. p values.

|  | sC2 vs sC1 | sC1 vs sC3 | sC3 vs sC4 | Mean error to identity |
| --- | --- | --- | --- | --- |
| T1 | 0,750 | 0,189 | 0,234 | 0.99% |
| T2 | 0,797 | 0,347 | 0,441 | 3.16% |

#### Segmentation pipelines comparison

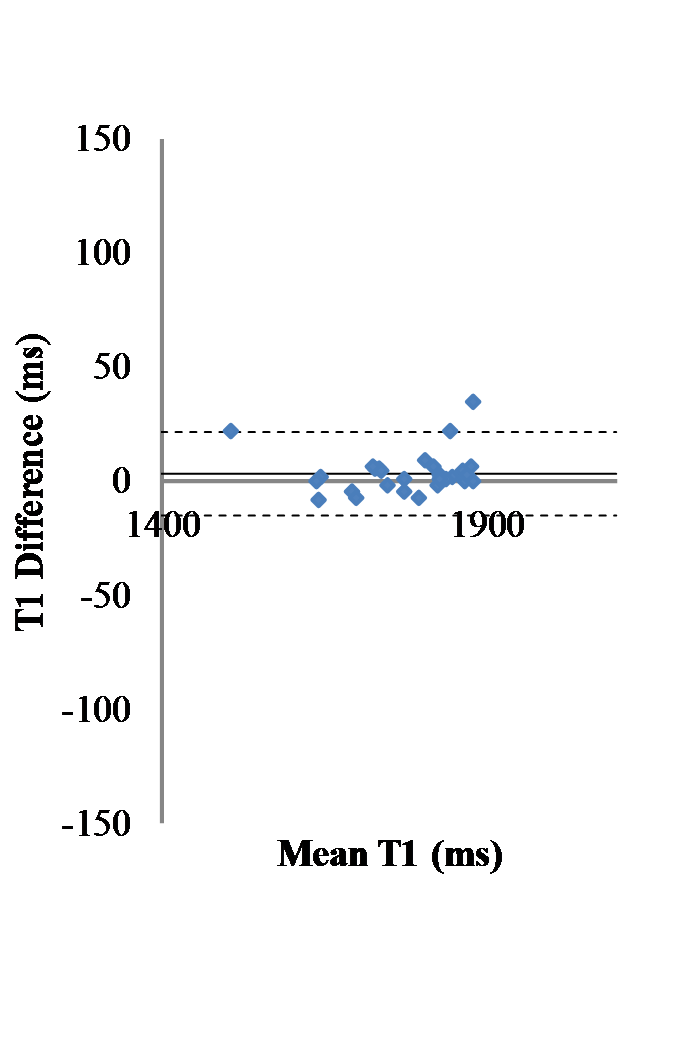

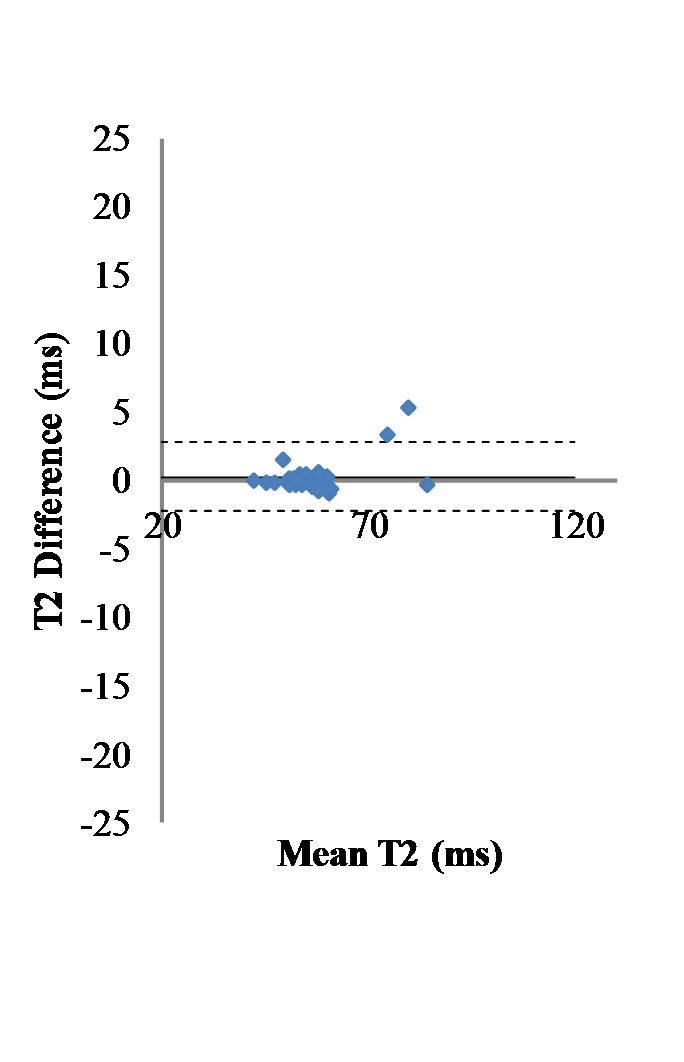

Figure 7: Comparison of segmentation pipelines. T1 (left) and T2 (right) relaxation times value differences measured for all regions of interest and average out of the whole set of animals (n=40) using the two different segmentation pipelines sC3 and sC4. fC2 fitting pipeline. Solid line: mean difference. Dashed line: ± two standard deviations.

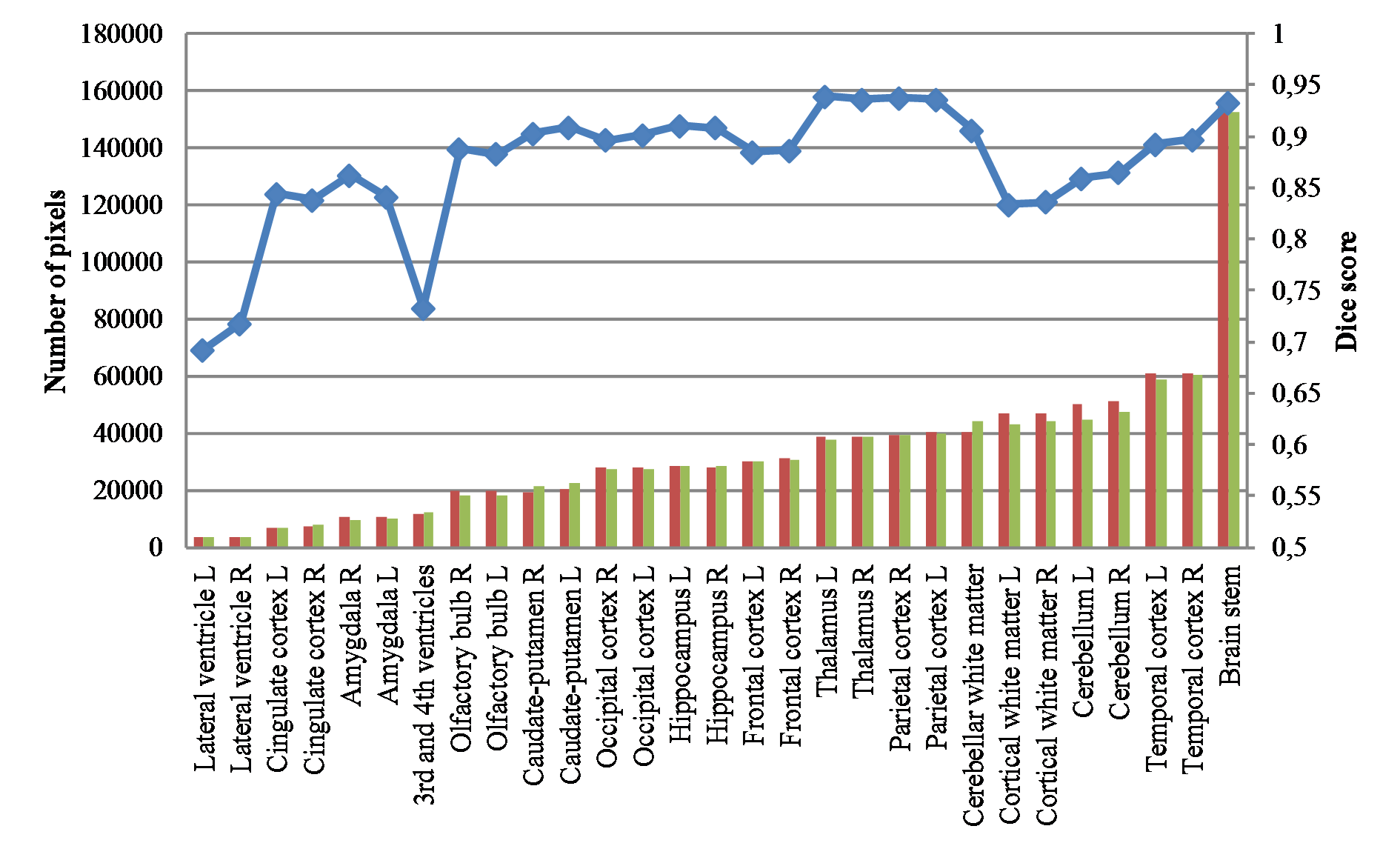

#### Figure 8: Differences between sC3 (Mircen) and sC4 (ICube) multi-atlas segmentation. Dice score (blue) and number of voxels using sC3 (green) or sC4 (red) in the different regions.
